# CAUTI’s Next Top Model – model dependent *Klebsiella* biofilm inhibition by bacteriophages and antimicrobials

**DOI:** 10.1101/2020.06.30.179804

**Authors:** Eleanor Townsend, John Moat, Eleanor Jameson

## Abstract

*Klebsiella* infections, including catheter associated urinary tract infections, are a considerable burden on health care systems. This is due to their difficulty to treat, caused by antimicrobial resistance and their ability to form biofilms. In this study, we investigated the use of a *Klebsiella* phage cocktail to reduce biofilm viability. We used two methodologies to investigate this, a standard 96-well plate assay and a more complicated Foley catheter-based model. The phage cocktail was used alone and in combination with clinically relevant antibiotic treatments. Viability was measured by both a resazurin based stain and colony forming unit counts, of cells sloughed off from the biofilm. We showed that phage infection dynamics and host survival vary significantly in different standard laboratory media, presumably due to the expression of different surface receptors and capsule composition by the bacteria effecting phage binding. This underscores the importance of a realistic model for developing phage therapy

We demonstrate that bacteriophage-based treatments are a viable option for preventing *Klebsiella* colonisation and biofilm formation on urinary catheters. Phage cocktails were able to significantly reduce the amount of biofilm that formed when they were present during early biofilm formation. The phages used in this study were unable to significantly reduce a pre-formed mature biofilm, despite encoding depolymerases. Phages applied together with antimicrobial treatments, showed synergistic interactions, in some cases the combined treatment was much more effective than antimicrobial treatments alone.

We show that phage cocktails have the potential to prevent *Klebsiella* biofilms in catheters, if used early or as a preventative treatment and will work well alongside standard antibiotics in the treatment of catheter-associated urinary tract infections (CAUTI).

## Introduction

Carbapenem-resistant and third-generation cephalosporin-resistant Enterobacteriaceae, such as *Klebsiella* sp. have been named by the World Health Organisation (WHO) as one of the critical priority bacteria in the fight against antibiotic resistance (Tacconelli et al. 2018). *Klebsiella* species cause a variety of opportunistic infections including urinary tract infection (UTI), pneumonia, septicaemia, wound infection, and infections in vulnerable patients including neonates and intensive care patients (Podschun and Ullmann 1998). *Klebsiella* species have a variety of virulence factors which make them efficient pathogens, but of most concern are the high levels of multiple antibiotic resistance mechanisms found within the genus. Resistance rates in *Klebsiella* have increased exponentially to most available antimicrobial drugs, and there have been cases of pan-resistant *Klebsiella* described (Sanchez et al. 2013; Elemam, Rahimian, and Mandell 2009). This increased resistance is often associated with an increased risk of mortality (Ben-David et al. 2012).

Complicating the antibiotic resistance of *Klebsiella* species is their ability to form biofilms. It has been shown with some antibiotics, that lack of penetration into the biofilm prevents killing of *Klebsiella*, however there are other mechanisms at work (Zahller and Stewart 2002). The thickness of mature *Klebsiella* biofilm has been implicated in increased antibiotic tolerance as well as biofilm heterogeneity (Singla, Harjai, and Chhibber 2014). Nutrient limitation, leading to lack of growth, has also been shown to contribute to antibiotic tolerance even with full penetration of the drug (Anderl, Franklin, and Stewart 2000; Singla, Harjai, and Chhibber 2014). It has also been shown that sublethal concentrations of antibiotics, often found within biofilms, can increase the virulence, biofilm formation and antibiotic resistance of *Klebsiella* species (Van Laar et al. 2015; Hennequin et al. 2012). *Klebsiella* strains that possess the ability to form rigid biofilms are also more likely to be Extended-Spectrum Beta-lactamase (ESBL) producers (Yang and Zhang 2008), as well as having increased ability to transfer plasmids within biofilm (Hennequin et al. 2012), which further complicates therapy.

The ability to form biofilms means *Klebsiella* spp. are able to colonise medical devices, leading to central line associated sepsis, ventilator associated pneumonia, and catheter associated urinary tract infections. *Klebsiella penumoniae* and *Klebsiella oxytoca* have both been found to be common causes of Catheter Associated Urinary Tract Infections (CAUTI) with high rates of antimicrobial resistance (Hidron et al. 2008; Wazait et al. 2003; Sabir et al. 2017).

Bacteriophage offer an alternative method of treating biofilm infections. The lytic lifecycle of phages is a virulent lifestyle that results in bacterial death, this lifestyle can be taken advantage of for therapeutic use. Utilising lytic phages as antibacterial agents was first described by d’Hérelle in 1919, and he summarised his successes in a review in 1931 (d’Herelle 1931). Phages offer a number of advantages over antibiotics as anti-biofilm agents, including increased specificity, accuracy and potency (Górski and Weber-Dabrowska 2005; Parisien et al. 2008). Phages are able to degrade the biofilm extracellular matrix, allowing antimicrobial penetration, lysis of bacterial cells and extensive biofilm disruption (Donlan 2001; Hughes, Sutherland, and Jones 1998).

Phage coating for catheters have already been developed for other common CAUTI pathogens such as *Proteus mirabilis, Escherichia coli, Pseudomonas aeruginosa* (Liao et al. 2012), and combinations of these bacteria in polymicrobial biofilms (Lehman and Donlan 2015; Carson, Gorman, and Gilmore 2010). Small scale clinical trials have been conducted, which show the efficacy of phage on catheters *in vivo* (Ujmajuridze et al. 2018). However, to our knowledge, there currently exists no equivalent treatment for *Klebsiella* caused CAUTI.

In this paper, we investigate the potential for a number of previously described *Klebsiella* phages (REF) to be used as anti-biofilm agents in preventing CAUTI. Using a number of urinary-isolated *Klebsiella* species we developed an *in vitro* model of CAUTI biofilms, which we used to test combinations of antimicrobial drugs alongside a phage cocktail. Our *in vitro* model was based around the use of artificial urine media (AUM) and sections of Foley catheter. We showed that the phage cocktail and antimicrobial therapy are in some cases able to complement each other’s activity, leading to an enhanced anti-biofilm effect.

## Materials and Methods

### Culture Conditions and Standardisation

Strains used in this study were from culture collections or clinical strains. The clinical strains (*Klebsiella pneumoniae* 170723, *Klebsiella oxytoca* 170748, *Klebsiella pneumoniae* 170958, and *Klebsiella oxytoca* 171266) were all isolated from urinary tract infections, with or without the presence of a medical device. Type strain *Klebsiella pneumoniae* subsp. *pneumoniae* 30104 was obtained from the DSMZ culture collection (Leibniz Institute, Germany). All isolates were stored long-term at −80°C and short-term at 4°C. Strains were maintained on LB agar and propagated in Cation-adjusted Mueller Hinton Broth (CAMHB). After growing to exponential phase, cells were rinsed in PBS and standardised by optical density at 600 nm before experimental work.

### Bacterial Growth Media

The bacterial growth media used within this study is specified for each technique, includes Cation-adjusted Mueller Hinton Broth (CAMHB, Sigma-Aldrich, Gillingham, UK), Luria Broth (LB, Sigma-Aldrich, Gillingham, UK), M9 minimal media (Elbing and Brent 2002) supplemented with 5mM glycerol and Artificial Urine Medium (AUM) (Brooks and Keevil 1997).

### Bacteriophage

Bacteriophage used in this paper were isolated and characterised by our lab group and described previously (Townsend et al. 2020). The phage used in this study, their isolation host strain, and source have been described in Table 1. Single phages were used for treatment, as well as a cocktail consisting of all six phages, which were mixed in equal portions, as determined by plaque forming units.

### Biofilm Formation in cell-culture plates

All *Klebsiella* strains were standardised to 1×10^2^ CFU/mL in CAMHB. Into a 96-well flat bottom cell culture treated plate, 200 µL of cell suspension was added. To form a biofilm, the plates were incubated statically for 16 hours at 37°C. All procedures were carried out in a Class II microbiological safety cabinet. Negative controls containing no inoculum, and positive controls which had no treatment applied, were included in each plate. All testing was carried out in triplicate, on three separate occasions.

### Phage and Antimicrobial Treatment in cell-culture plates

For investigating phage inhibition of biofilm formation, 10 µL of phage suspension was added to the well before any bacteria were added. Biofilms were then allowed to form, as described above.

To treat mature biofilms, the biofilms were first rinsed in PBS to remove any planktonic cells. Following this, 10 µL of phage stock suspension (1×10^2^ PFU/mL) was added to each well. For treatment with meropenem, the drug was suspended in CAMHB at 128 mg/L and 64 mg/L for high and low concentrations respectively. Untreated controls and negative controls were included in every experiment. The experiments were performed in triplicate.

### Phage infection in different media

*Klebsiella* strains were grown to exponential phase in four different media (LB, CAMHB, M9 supplemented with 5mM glycerol and AUM (Brooks and Keevil 1997)). The strains were then diluted to an OD_600nm_ which was equivalent to approximately 1 × 10^7^ CFU/mL and 150 µL of bacterial suspension was added per well in a 96-well microtitre plate. The wells then had either 50 µL phage cocktail (test wells) or 50 µL media (control wells) added.

The 96-well microtitre plate was sealed and incubated at 37°C with shaking in a FLUOstar® Omega plate reader (BMG Labtech, Aylesbury, UK). The optical density of the cultures was recorded over 24 hours, with readings at OD_600nm_ taken every 5 minutes. Each plate contained technical duplicates for each *Klebsiella* strain in each media and was performed in biological triplicate.

The area under the curve (AUC) was calculated using MatLab. The ratio of bacteria-only control AUC to phage infected AUC was calculated for each growth media. For each strain, the AUC ratio was normalised by Ln transformation and then each media was compared using a two-tailed t-test to identify where phage infection was affected by the growth media.

### Biofilm formation and treatment in simple catheter model

Foley catheters (Folatex ref. AA1B16 Ch/Fr 16/5.33, 30-45 ml/cc, silicone coated latex urinary catheter/straight/2-way) were cut into 1.5 cm long sections, aseptically. The catheter sections were incubated in a bacterial suspension (1×10^2^ CFU/mL in CAMHB) for 2 hours, at 37°C at 150 rpm. Following incubation, foley catheters sections were transferred to a 24-well cell-culture plate (BRAND) containing 1.5 mL AUM (Brooks and Keevil 1997). Phage cocktail (1×10^2^ PFU/mL) was added at the initiation of biofilm formation, where it was diluted 1 in 7 in 1.5 mL AUM. Biofilms with and without phage cocktail were incubated statically for 16 hours at 37°C. Before antimicrobial treatment, catheter sections were rinsed in PBS to remove any planktonic or loosely adhered cells. Meropenem, mecillinam, or trimethoprim were then added at 164 mg/L or 64 mg/L in AUM to the biofilm. As trimethoprim was dissolved in Dimethyl sulfoxide (DMSO), as an additional control, DMSO was added to AUM without trimethoprim. Meropenem and mecillinam stocks were dissolved in water. After the addition of drug, biofilms were further incubated for 5 hours before being removed for analysis. Untreated controls and negative controls were included in every experiment. The experiments were performed in triplicate.

### Biofilm Viability Analysis

Media and treatments were removed from the biofilm by pipetting off overlying media followed by rinsing in PBS, and viability was assessed using 10 µg/mL resazurin sodium salt. Absorbance was measured at OD _570 nm_ and _600nm_ and used to calculate the percentage viability, relative to negative controls incubated in each plate (Kirchner et al. 2012).

For the Foley Catheter biofilms, the viability was also analysed by incubating the treated section of catheter in 2 mL CAMHB for 2 hours at 37°C at 150 rpm. This regrowth bacterial suspension was then used for Miles and Misra counts to assess the CFU/mL. Briefly, the bacterial suspension was serially diluted 1 in 10 and then 10μL plated out on LB agar plates in triplicate. The plates were then incubated overnight at 37°C, the colonies were counted and used to calculate the bacterial concentration in original suspension.

### Statistical Analysis

Graph production was carried out in MatLab (MatLab R2020a) and GraphPad Prism (Prism version 8.2.4). Unpaired two-tailed t-tests were used to establish significant differences between treated samples and untreated controls. Percentage viability scores were log_2_ transformed before statistical analysis took place. A 2-way ANOVA with Sidak’s multiple comparison test was used to establish any differences between phage infection in different bacterial growth media. Statistical significance was achieved if p < 0.05.

## Results

### Solo phage treatment is able to prevent biofilm formation, but the phage cocktail has a wider range of activity

Phage had been added to the biofilm grown in CAMHB in 96-well microtitre plates, both at the initiation of biofilm formation (Figure 2A), and after biofilms had matured, for a treatment period of 5 hours (Figure 2B).

**Figure 1.**
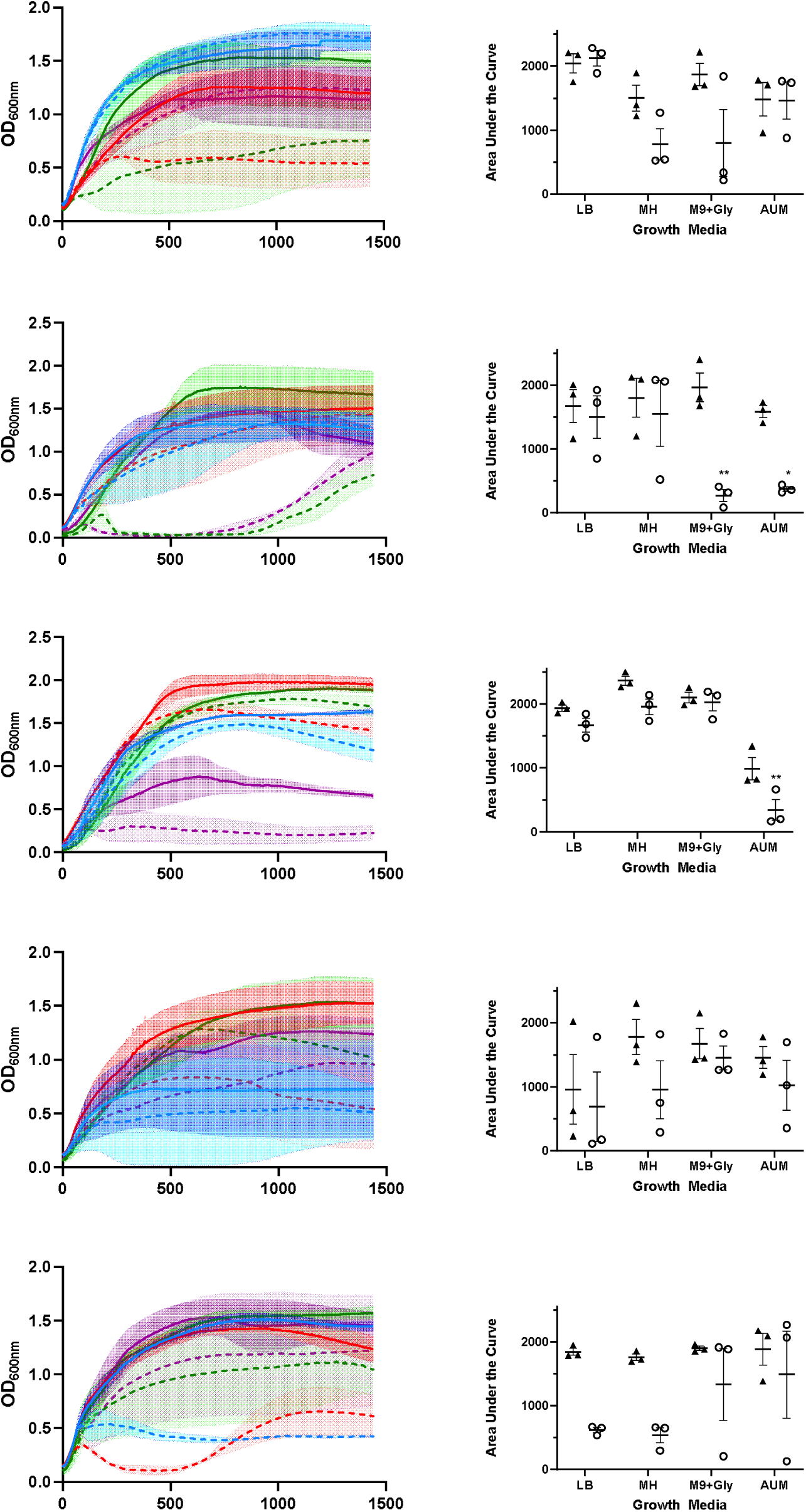
Infection curves and area under the curve analysis for five *Klebsiella* strains infected with the same phage cocktail in four different media. Infection of five *Klebsiella* strains was performed as described in the methods and monitored spectrophotometrically over 24 hours (**A**). Bacteria only controls were performed alongside bacteria infected with phage cocktail. Line displays the mean optical density, error bars display standard error across biological triplicates. **B**. Area under the curve analysis was performed on the curves displayed in figure 1a.* Denotes significant difference (p<0.05) between bacterial growth with and without phage cocktail.

**Figure 2.**
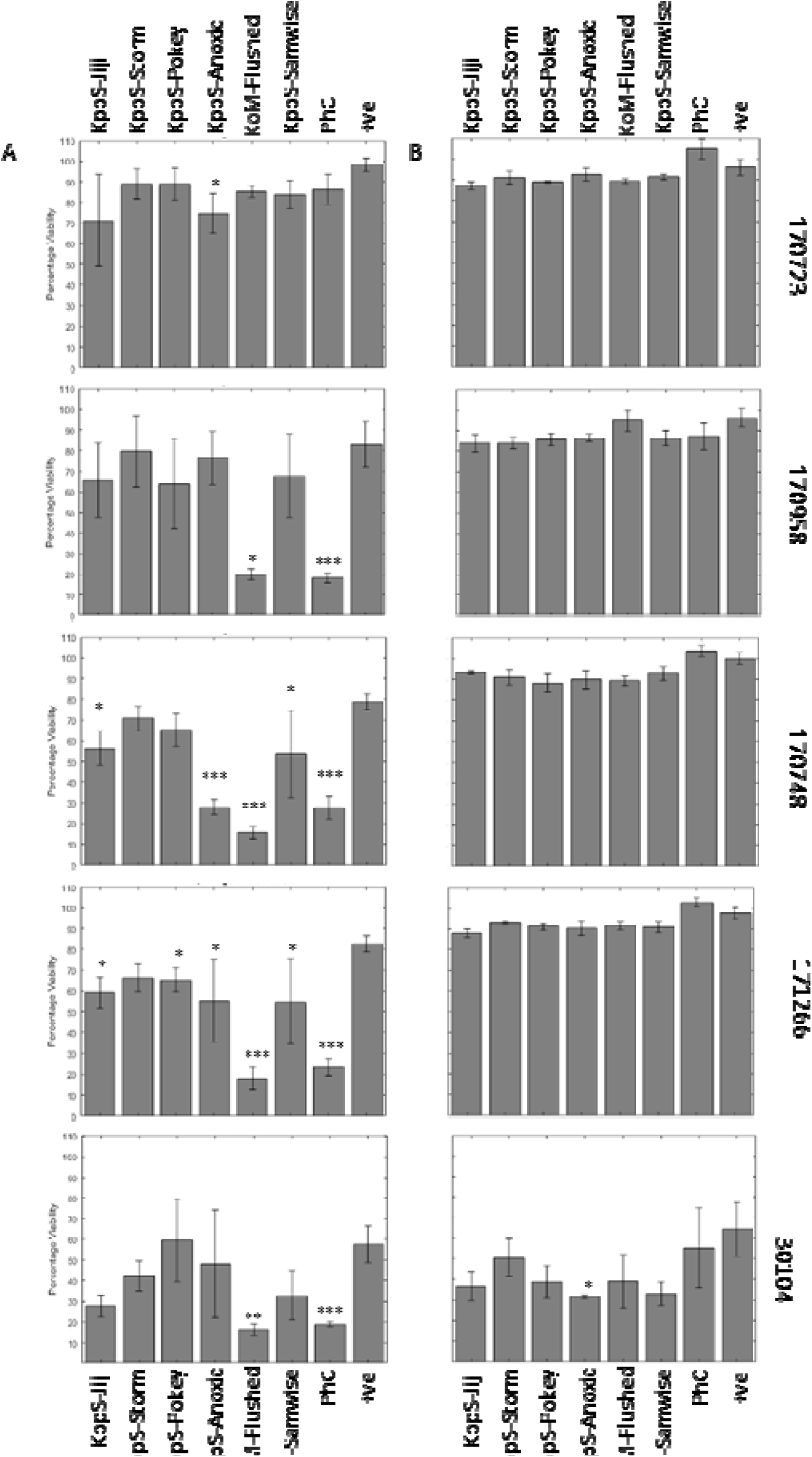
Individual phage and phage cocktail are able to reduce the viability of some, but not all, *Klebsiella* biofilms. Phage were added to biofilms either individually or combined in a cocktail (PhC) at the start (A) of biofilm formation or for five hours (B) to a mature *Klebsiella* biofilms, as described in the methods. Some individual phage were able to reduce biofilm viability when added at the start of biofilm formation (A), and in four strains the cocktail was effective when added at this point. When used on a mature biofilm (B), there was very limited effects on biofilm viability. Symbols denote significant difference (t-test); * compared to positive control.

Solo phage treatment was able to prevent biofilm formation in a number of strains (Figure 2A), although strains varied in susceptibility to individual phage. Phage KoM-Flushed had the widest host range, being able to significantly reduce biofilm formation in four out of five strains (170748, 170958, 171266, and 30104). Phage KppS-Anoxic was able to significantly reduce biofilm viability in three strains (170748, 170723, 171266), while phages KppS-Samwise and KppS-Jiji both caused significant reductions in 171266 and 170748. Phage KppS-Pokey was only able to significantly reduce biofilm formation in 171266. Finally, phage KppS-Storm was unable to cause any significant reductions in biofilm formation in any strains. When these phages were combined to create a phage cocktail, this treatment significantly reduced biofilm formation in all strains except 170723.

When added onto a mature biofilm (figure 2B) there were no significant reductions in the viability of the biofilm, with the exception of phage KppS-Anoxic in strain 30104. Therefore, at this point the results already showed that phage treatment is more effective at preventing biofilm formation rather than penetrating and killing a mature biofilm community.

### Combination of the phage cocktail and meropenem is more effective than monotreatments

Biofilms were again grown in CAMHB in 96-well microtitre plates and challenged with two different doses of meropenem, the phage cocktail, and a combination of meropenem and phage cocktails (Figure 3). The phage cocktail was again added at the start of biofilm formation (Figure 3A) or onto a mature biofilm (Figure 3B), while meropenem treatments were added after the biofilm had matured.

**Figure 3.**
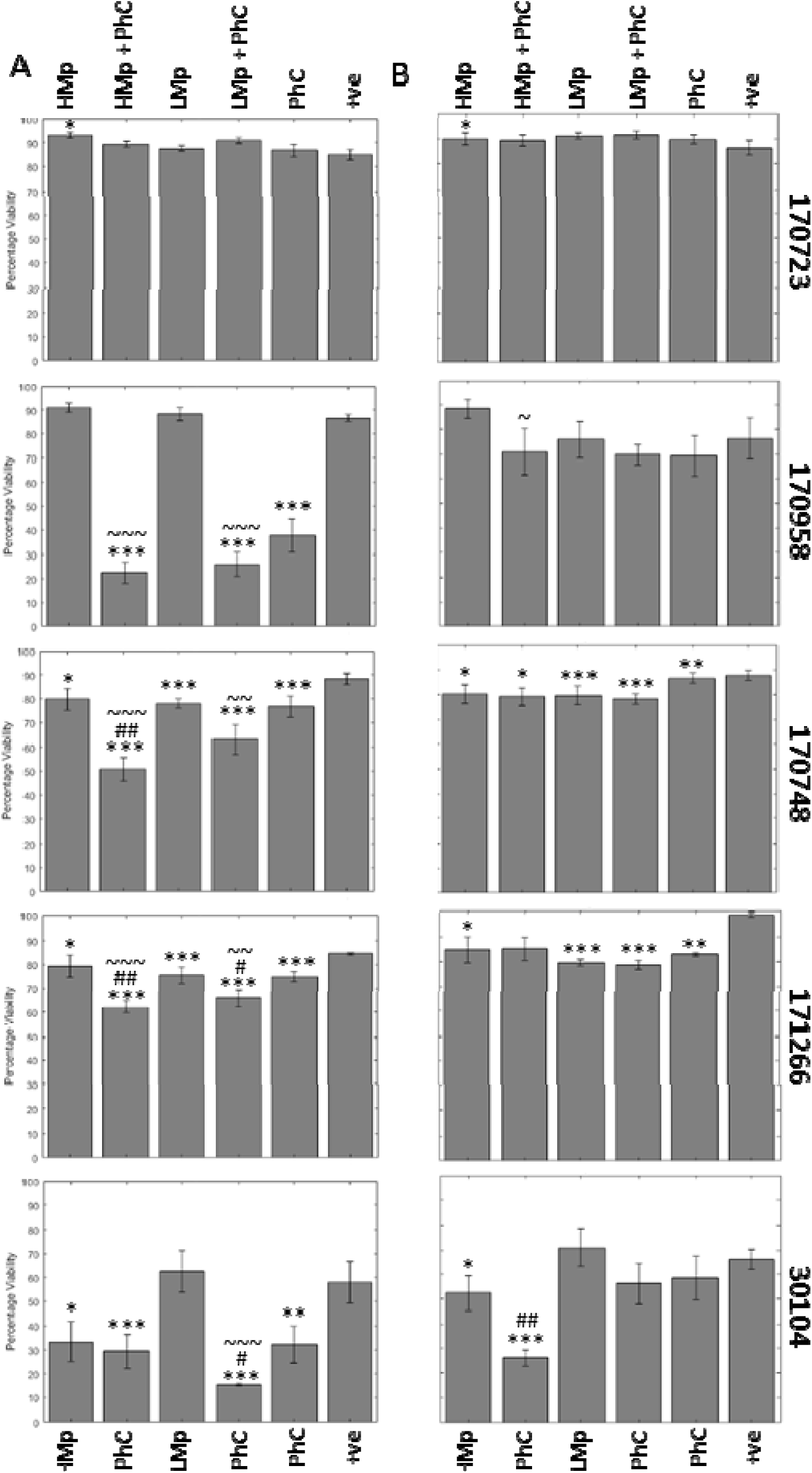
There is synergy between phage cocktail and meropenem when used to prevent biofilm formation. Phage treatments when added at the start (A) or for five hours (B) to the *Klebsiella* biofilms as described in the methods. *Klebsiella* strain 170723 had no significant reductions in biofilm viability, however all other strains showed some reductions, and there appeared to be a synergistic effect when used in combination with meropenem. Symbols denote significant difference (t-test); * compared to positive control, # compared to corresponding phage cocktail treatment, and ∼ compared to corresponding meropenem treatment.

As in previous tests, 170723 was unaffected by the phage cocktail, as well as the meropenem treatment at both time points. In fact, there was a small significant increase in viability when the biofilm was exposed to high levels of meropenem.

In strain 170958, dramatic effects were seen when the phage cocktail was added at the start of biofilm formation, but these were not seen when the phage was added to a mature biofilm. There was an additive effect between meropenem and the phage cocktail when added at the start, where the combination caused a further reduction in viability compared to the treatment with meropenem alone, but not the phage cocktail alone.

For *Klebsiella* strain 170748, there was a complementary effect shown between the antimicrobial and the phage cocktail, where a combination of the two treatments significantly reduced the biofilm compared to either treatment alone. When the phage cocktail alone was added at the start, it was able to cause a significant decrease in the viability of the biofilm, and this reduction was more dramatic when in combination with the meropenem treatment. When added to the mature biofilm, the treatments were able to cause a reduction in the viability of the biofilm, but the complementary effect was no longer evident.

Strain 171266 was the most susceptible to the two treatments in the 96 well plate assay. When the phage cocktail was added at the start, all treatments were able to cause significant reductions in the biofilm. This was especially evident in those treated with both the phage cocktail and meropenem, there was again evidence of complementarity between meropenem and the phage cocktail. When treating the mature biofilm, while there were some significant reductions in the biofilm viability, there was no longer evidence of synergistic activity.

The type strain 30104 also showed complementarity between the antimicrobial and the phage cocktail was also moderately susceptible to treatment but does not form as robust a biofilm in comparison to the other strains. However, the phage cocktail alone was still able to cause a significant reduction. On the mature biofilm, these effects were less evident, and the phage cocktail was not effective.

### Phage infection varies dependent on growth media

To assist in the development of a more reflective *in vitro* model of CAUTI, we first investigated phage infection in a series of rich and more minimal growth media. We continued with our use of CAMHB, and additionally tested in LB, M9 supplemented with glycerol as a carbon source, and AUM. We suspected that CAMHB as a rich media would not be very realistic, but had concerns if phage infection would still occur in more minimal and restrictive growth media.

The phage cocktail was used to infect each bacterial strain planktonically in four different growth media, LB, CAMHB, M9 supplemented with 5mM glycerol, and AUM (Figure 1A). As rates of bacterial growth also differ in each media, the ratio of area under the curve (AUC) in a bacteria only control and bacteria infected with phage was calculated in each media (Figure1B).

An ANOVA with Sidak’s multiple comparison tests was used to analyse the data. The data was analysed for interaction effects, and in all but 170958, the two factors (phage and media) did not interact. Therefore, in the four remaining strains, the effects of phage and media were independent of each other.

In strain 171266, neither the effects of media or phage were considered significant, nor were there any significant differences within each media in the presence or absence of phage. However, in each media there is a slight decrease in the mean AUC in the presence of phage.

For the type strain 30104, there is no effect of the media on the variance, but the presence or absence of phage accounts for 37.12% of the variance and is considered significant (p value = 0.0021). There were no significant differences between +/− phage in each media, but it can be seen that there was a more marked decrease in AUC in the LB and MH media compared to M9/Gly and AUM.

Both phage and media account for variance in the data in strain 170723 (12.44% and 32.95%, respectively) and this is also significant (p value = 0.0390, p value = 0.0185, respectively). It can be observed that there is little reduction of the AUC in the LB and AUM media, but in MH and M9/Gly there is a decrease in AUC.

In strain 170748, again both phage and media account of a significant portion of the variance (p value = 0.0008, 7.145% variance, and p value < 0.0001, 83.51% variance, respectively). While there were modest decreases in AUC with phage in media LB and MH, only the decrease in AUM media was significant (0.0059).

The results in 170958 are difficult to interpret as there is interaction of the two factors (p value = 0.0349, 20.07% variance). The effect of phage is considered significant (p value = 0.0006, 33.5% variance), while the effect of media is near significant (p value = 0.0542, 17.23% variance). There was also a significant decrease in AUC with phage in both M9/Gly (0.0019) and AUM (0.0262), while there was very little change in LB and MH.

Overall, these results show that the nutritional environment that the host bacteria are growing in can influence the cell lysis by phage, although this varies dependent on the host strain. Therefore, we decided to continue our experiments using AUM for a growth media, as this both allowed phage infection to occur in a number of strains planktonically, as well as being close to the clinical situation we are attempting to mimic.

### Biofilms in the *in vitro* Foley catheter model

The phage cocktail was then tested in an *in vitro* Foley catheter model using AUM, as described in the methods. Phage treatment was added at the start of the biofilm formation and then antimicrobials were added later to the mature biofilm. Biofilms in the Foley catheter model were analysed in two ways, viability staining and CFU counts from the dispersal of the biofilm, as described in the methods.

Continuing with meropenem treatment, in three strains (170723, 170958, and 170748), the combination treatment of the phage cocktail and meropenem significantly reduced the viability of the biofilm (Figure 4A). For 170723, this was significantly lower than both the drug alone and the phage cocktail alone, meaning there is a complementary effect between the two treatments. For strains 171266 and 30104, there were reductions in the viability of the biofilm in response to both treatments, alone and combined, but these were not significant.

**Figure 4.**
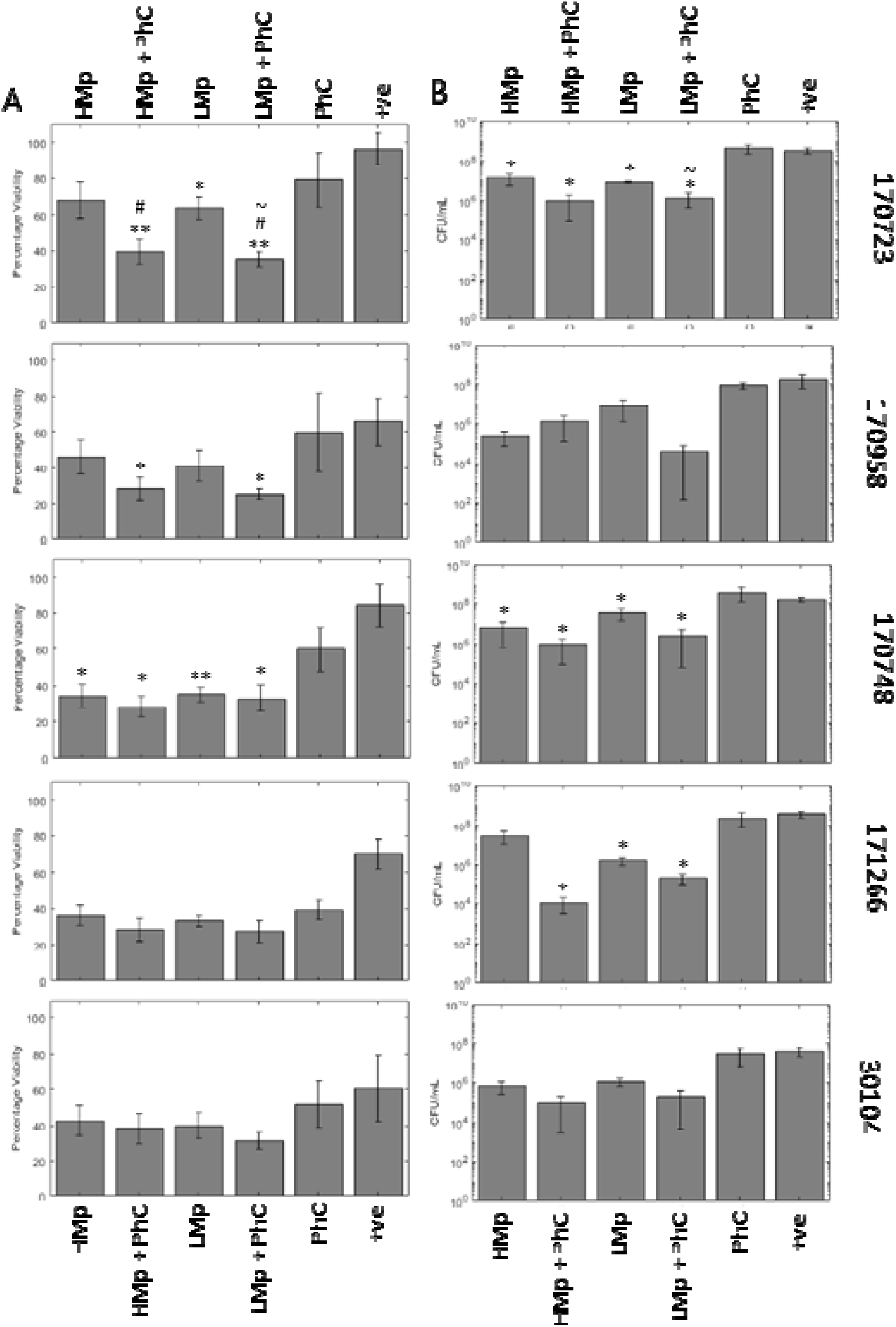
In a more complex model (Foley catheter *in vitro* model) treatments are less effective but are still able to cause some reduction in biofilm viability and sloughing. Phage treatment was combined with meropenem in a Foley catheter *in vitro* model. Biofilm reduction was measured by viability assay (A) and CFU counts (B). Treatments were able to reduce viability and sloughing, most notably in strain 170723 which had previously not responded to the same treatment in previous testing. Symbols denote significant difference (t-test); * compared to positive control, # compared to corresponding phage cocktail treatment, and ∼ compared to corresponding meropenem treatment.

The second analysis was of colony forming units counts of bacterial cells sloughed off from the biofilm (Figure 4B). This should be proportional to what is within the biofilm itself, as well as giving an indication to likelihood of infection to disperse from the biofilm and disseminate after treatment. Here the results are less clear, with only 170723, 170748, and 171266 showing significant reductions in viability after treatment with either meropenem or combination. Only 170723 treated with low level meropenem and the phage cocktail showed a significant reduction compared to low level meropenem alone.

The next drug used was mecillinam, a second-line antibiotic used for CAUTI (NICE 2018). *Klebsiella* should be intrinsically resistant to this penicillin antibiotic, therefore it is not surprising that the phage treatment was driving the reduction in viability of the biofilm here (Figure 5A). Phage treatment often resulted in the lowest *Klebsiella* viability, lower than that of the phage cocktail combined with mecillinam. In strains 170723, 170958, and 171266, the combination treatments were significantly lower viability compared to the mecillinam only treatments. The colony forming unit analysis showed no significant differences regardless of treatment (Figure 5B).

**Figure 5.**
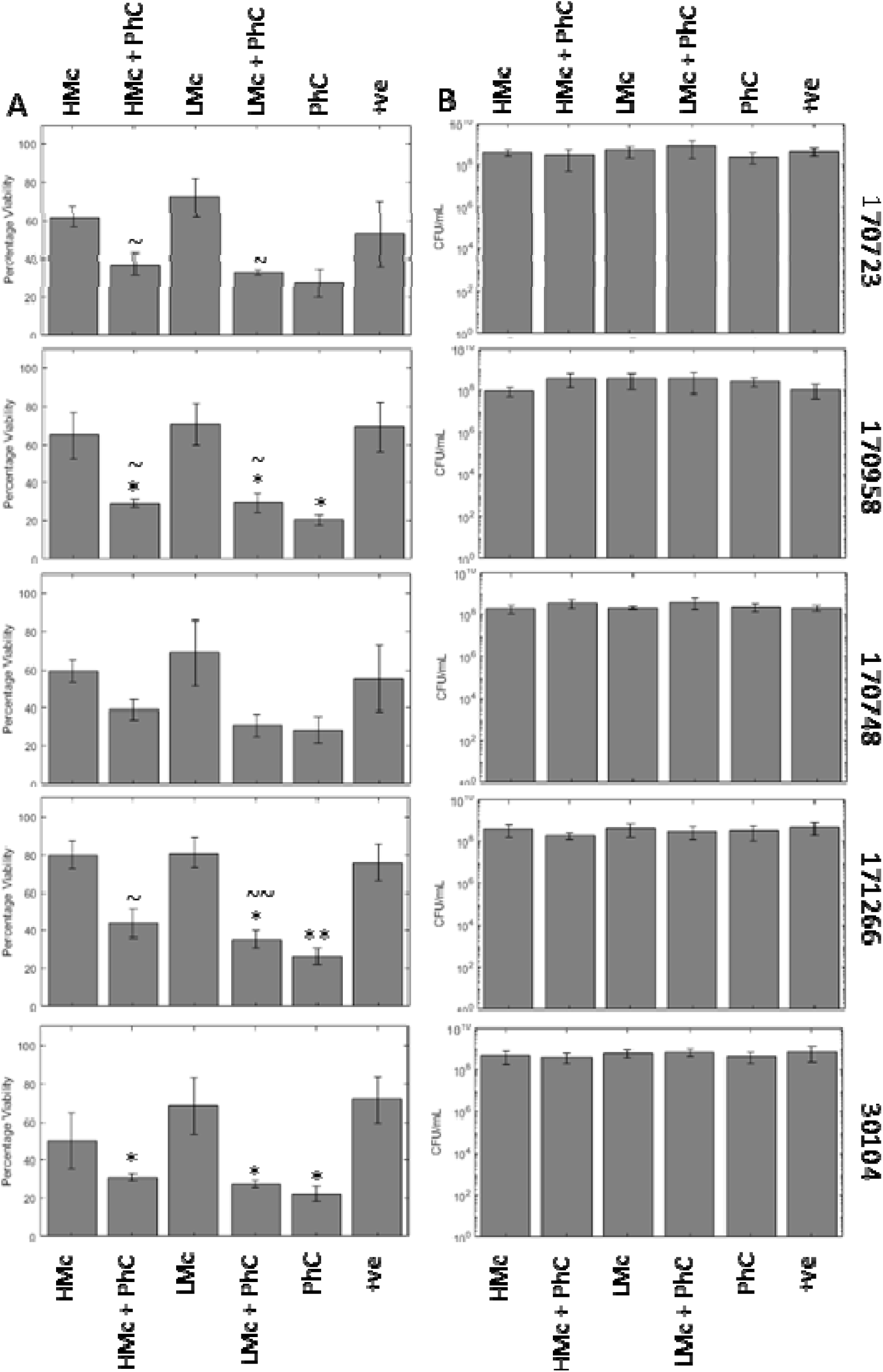
Mecillinam was not effective at reducing *Klebsiella* biofilms by either measure, and phage cocktail was more effective alone. Phage treatments were combined with mecillinam, a second line drug used for treatment of CAUTI in the *in vitro* Foley catheter model. The treatment effect was measured by viability assay (A) and CFU counts (B) as described in the methods. The viability staining showed that phage cocktail alone was more effective compared to drug alone or in combination. All effects were lost when measured by CFU counts. Symbols denote significant difference (t-test); * compared to positive control, # compared to corresponding phage cocktail treatment, and ∼ compared to corresponding mecillinam treatment.

Trimethoprim was the final drug used, a first-line antibiotics in the treatment of CAUTI (NICE 2018). As the drug needed to be suspended in DMSO, all other treatments and the positive control also had DMSO added to control for any additional effects.

In strain 170958 and 30104, all treatments significantly decreased the percentage viability of the biofilms. In 170748, high trimethoprim with the phage cocktail, low trimethoprim, and low trimethoprim with phage significantly reduced the viability of the biofilm (Figure 6A). In strain 171266, all treatments except low trimethoprim were able to significantly reduce the viability of the biofilm. In strain 170723, despite reductions in viability, none of the changes caused by the treatments were statistically significant. However, when analysed by CFU counts, this change became significant (Figure 6B). Strain 170748 also had significantly lower CFU counts with every treatment, with the addition of high trimethoprim and the phage cocktail being a more significant reduction compared to trimethoprim alone. In strains 171266 and 30104, the combined treatments significantly reduced the CFU counts compared to the phage treatment alone. Strain 170958 had no significant changes to the CFU counts.

**Figure 6.**
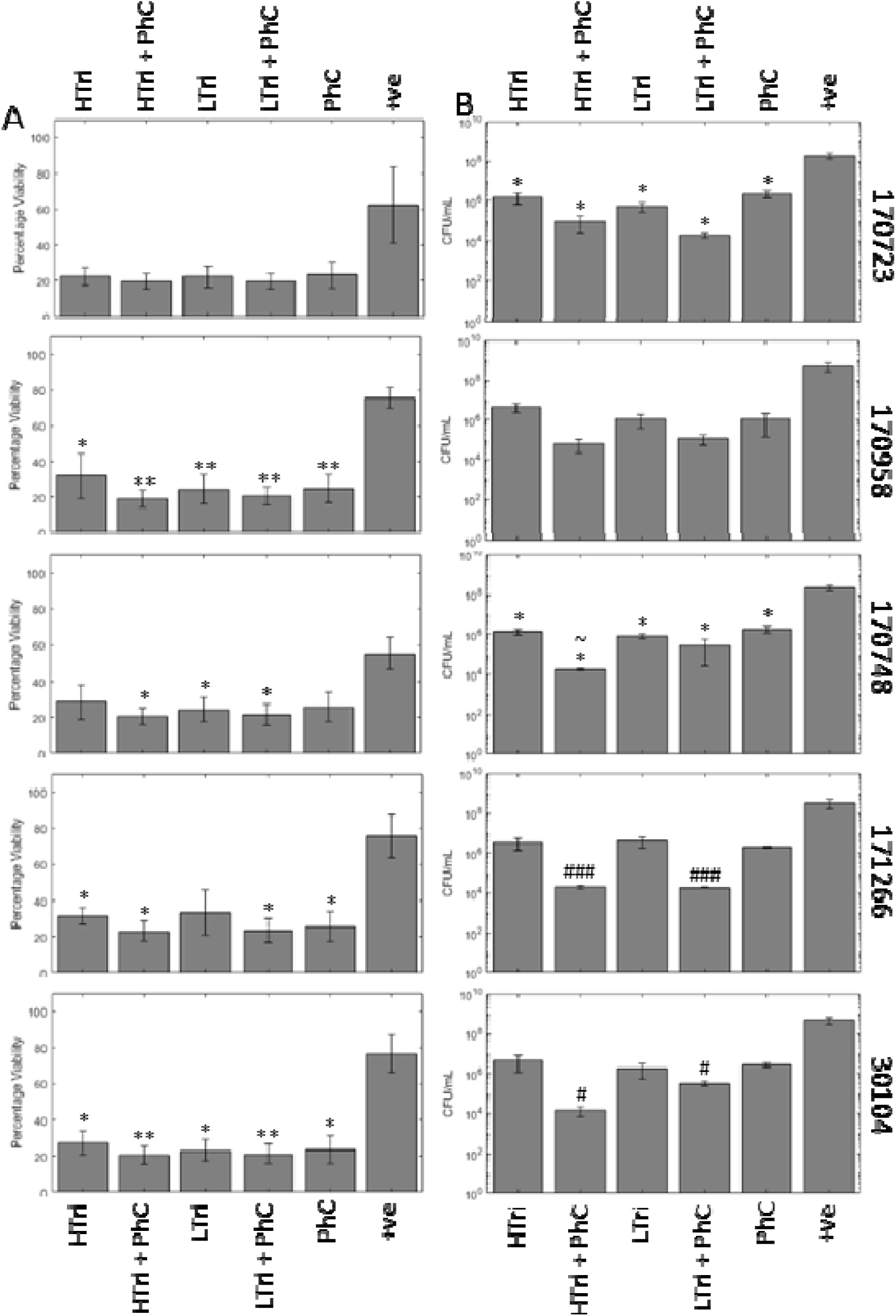
Trimethoprim is effective at reducing *Klebsiella* biofilms and CFU counts show it has some additive action with phage cocktail. Phage treatments combined with trimethoprim, a first line drug for CAUTI, measured by viability assay (A) and CFU counts (B) in an *in vitro* Foley catheter model. In the viability assay, all treatments were approaching the lower limit of detection of the assay, therefore it is difficult to see any additive or synergistic effect of the phage treatment. However, this is more evident in the CFU counts where reductions are greater in the combination treatments. Symbols denote significant difference (t-test); * compared to positive control, # compared to corresponding phage cocktail treatment, and ∼ compared to corresponding trimethoprim treatment.

## Discussion

In this paper we have shown that antibiotics can be used in combination with phage cocktail treatments, synergistically to reduce the amount of biofilm formation on urinary tract catheters. This shows the potential of using phages to prevent *Klebsiella* CAUTI and the further complications associated with *Klebsiella* infections.

We have also shown that using appropriate models that replicate the *in vivo* conditions as accurately as possible is important. This has been shown to be important for antibiotic treatments, but even more so for phage infection, evidenced by the initial experiments comparing media for planktonic phage infection and the varying response of strain 170723 in the two main biofilm models. Although the model presented here remains relatively simple, we have already shown that increasing complexity is important in testing new treatments. We theorise that on exposure to different media and surfaces, *Klebsiella* may vary their capsule composition and cell surface receptors in the different environments and nutrient availability (Ratner, Sampson, and Weiss 2015; Huang et al. 2012; Stewart and Olson 1992; Cano et al. 2015).

Mixtures of phages, commonly known as phage cocktails, are used for therapeutic reasons over a lone phage for the same reason that drug cocktails are often used in particularly recalcitrant infections such as HIV or TB. Phage cocktails have a broader host range than individual phages, reducing the chance of phage resistance emerging, which can maximise the efficiency of therapy (Kutateladze and Adamia 2010). Phage resistance is a common phenomenon in the natural predator prey relationship between phages and bacteria. More specifically in the case of *Klebsiella* biofilms, phages may possess depolymerase enzymes (Wu et al. 2019; Geredew Kifelew, Mitchell, and Speck 2019) that are able to degrade either the bacterial capsule or the extracellular matrix that protects cells within the biofilm. Depolymerase action can then facilitate the penetration of antimicrobial drugs into the biofilm or may uncover receptors for a secondary phage infection (Verma, Harjai, and Chhibber 2010). In this way, phages are protected from resistance developing and the likelihood of synergistic interactions increases. A phage cocktail was composed of six phages that had been previously characterised by our group (Townsend et al. 2020). In this study, we showed that in a number of cases, phages combined with an antimicrobial drug were able to cause a significant reduction in biofilm viability compared to each treatment alone. We theorise that the treatments have a synergistic action when used together, but the precise mechanism behind is yet to be elucidated. To further characterise this complementary effect, investigations into the pharmacokinetics of the treatments would need to be established for calculating synergistic effects of combining phage and antimicrobials.

Our study focusses mainly on the use of meropenem, a carbapenem antibiotic. In addition, we have used trimethoprim, a sulphonamide, and mecillinam, a penicillin, which are both recommended in the NICE guidelines for use in treatment of CAUTI (NICE 2018). Trimethoprim was particularly effective against the *Klebsiella* strains used in this study. So much so, that in the viability assay the results were approaching the lower limit of detection of the assay, making it difficult to detect any additional benefit of co-treatment with phage. However, these effects were better seen in the colony forming units counts of cells sloughed off the biofilm, highlighting the benefits of using multiple methods to measure treatment effects. Conversely, mecillinam was not at all effective against the strains. This is not surprisingly as *Klebsiella* have long been recognised as intrinsically resistant to penicillin antibiotics (Hamilton-Miller 1965). The hypothesis behind choosing mecillinam was that there may be some co-operative activity between the phage and drug which would overcome this resistance. This has been previously demonstrated where a phage’s depolymerase enzymes are able to degrade the matrix and capsule of a bacteria, allowing antibiotic penetration where it would not normally occur (Tait, Skillman, and Sutherland 2002; Bansal, Harjai, and Chhibber 2014). Unfortunately, this was not the case here. It is well characterised that sub-inhibitory concentrations of antibiotics can stimulate biofilm formation (Dunne 1990). Instead this appears to be the effect seen here, where the phage cocktail is the driving force in the reduction of viability and the presence of mecillinam only “irritates” the biofilm (Van Laar et al. 2015; Hennequin et al. 2012). While the use of a phage cocktail can theoretically assist drug penetration and increase efficacy, it cannot substitute selection of an appropriate antibiotics.

In the development of our *in vitro* CAUTI model, we investigated phage infection in different nutritional environments. The effects of nutritional environment were briefly investigated by Storms et al. in the development of the virulence index, a method of quantifying phage virulence relative to the natural host growth rate (Storms et al. 2019). In this study it was also shown that temperature had an effect on phage virulence alongside growth media, as well as the same phage having different dynamics in secondary host strains. These are all factors that need to be taken into consideration when optimising phage cocktails for use outside of the lab. In this work, all experiments were carried out at 37°C to mimic the temperature experienced in human infections.

There have been a number of studies for other urinary tract pathogens, where phages have been shown to be successful prevention strategies (Milo et al. 2017; Nzakizwanayo et al. 2016; Lehman and Donlan 2015; Carson, Gorman, and Gilmore 2010), sometimes in combination with a benign biofilm (Liao et al. 2012). These results have been shown in clinical or urinary tract pathogens (but not *Klebsiella*) without any negative side effects being found to be associated with the phage treatment (Ujmajuridze et al. 2018). Although larger studies are required before phage treatments can overtake antibiotics, the current evidence shows phage treatments are accepted by patients and can show efficacy. A review of current phage therapeutics, including urinary tract catheters can be found here (Ryan et al. 2011). We propose that something similar could be developed for *Klebsiella*. Phages have already been shown to be a valid method of eradicating *Klebsiella* biofilms, even in multi-resistant strains (Jamala et al. 2015; Bedi, Verma, and Chhibber 2009; AlBany, Al-Berfkani, and Assaf 2019), depolymerases isolated from phage have also shown to have efficacy (Majkowska-Skrobek et al. 2016). Our Klebsiella model shows that phage cocktails have promise in the prevention of *Klebsiella* CAUTIs, which would prevent complications in vulnerable patients and prolong the time urinary catheters can remain *in situ*.

## Conclusions

This is the first time our *in vitro* CAUTI model has been presented, and the first time the efficacy of antibiotics in combination with phage cocktail has been tested on *Klebsiella* CAUTI biofilms. Phage cocktails shows synergy with the current CAUTI antimicrobial treatments, trimethoprim and meropenem, to help prevent biofilm formation by *Klebsiella* species in catheters. This has important implications for clinical therapy, to offer an alternative treatment for antimicrobial resistance *Klebsiella*.

## Acknowledgements

WISB is a BBSRC/EPSRC Synthetic Biology Research Centre (grant ref: BB/M017982/1) funded under the UK Research Councils’ Synthetic Biology for Growth programme.

---

**Table 1.** Description of the phages used in this work. Phages used in this work are described within this table including their full name, isolation host and source of isolation.

| Full Name | Isolation host | Source |
| --- | --- | --- |
| vB_KppS-Samwise | Klebsiella pneumoniae subsp. pneumoniae 30104 | Slurry |
| vB_KppS-Jiji |  | Pond water |
| vB_KppS-Storm |  | Storm tank storage |
| vB_KppS-Pokey |  | Anoxic sludge |
| vB_KppS-Anoxic |  | Anoxic sludge |
| vB_KoM-Flushed | Klebsiella pneumoniae 170958 | Sewage – mixed liquor |

## References

AlBany, Yousif Abdullah, Mohammad Ismail Al-Berfkani, and Mahde Saleh Assaf. 2019. ‘Phage Therapy Against Biofilm of Multidrug-Resistant Klebsiella Pneumoniae Isolated from Zakho Hospital Samples’, Polytechnic Journal, 9: 17–22.

Anderl, Jeff N, Michael J Franklin, and Philip S Stewart. 2000. ‘Role of antibiotic penetration limitation in Klebsiella pneumoniae biofilm resistance to ampicillin and ciprofloxacin’, Antimicrobial Agents and Chemotherapy, 44: 1818–24.

Bansal, Shruti, Kusum Harjai, and Sanjay Chhibber. 2014. ‘Depolymerase improves gentamicin efficacy during Klebsiella pneumoniae induced murine infection’, BMC Infectious Diseases, 14: 456.

Bedi, Manmeet Sakshi, Vivek Verma, and Sanjay Chhibber. 2009. ‘Amoxicillin and specific bacteriophage can be used together for eradication of biofilm of Klebsiella pneumoniae B5055’, World Journal of Microbiology and Biotechnology, 25: 1145.

Ben-David, D, R Kordevani, N Keller, I Tal, A Marzel, O Gal-Mor, Y Maor, and G Rahav. 2012. ‘Outcome of carbapenem resistant Klebsiella pneumoniae bloodstream infections’, Clinical Microbiology and Infection, 18: 54–60.

Brooks, T, and CW Keevil. 1997. ‘A simple artificial urine for the growth of urinary pathogens’, Letters in applied microbiology, 24: 203–06.

Cano, Victoria, Catalina March, Jose Luis Insua, Nacho Aguiló, Enrique Llobet, David Moranta, Verónica Regueiro, Gerard P. Brennan, Maria Isabel Millán-Lou, and Carlos Martín. 2015. ‘Klebsiella pneumoniae survives within macrophages by avoiding delivery to lysosomes’, Cellular microbiology, 17: 1537–60.

Carson, Louise, Sean P. Gorman, and Brendan F. Gilmore. 2010. ‘The use of lytic bacteriophages in the prevention and eradication of biofilms of Proteus mirabilis and Escherichia coli’, FEMS Immunology & Medical Microbiology, 59: 447–55.

d’Herelle, F. 1931. ‘Bacteriophage as a Treatment in Acute Medical and Surgical Infections’, Bulletin of the New York Academy of Medicine, 7: 329–48.

Donlan, Rodney M. 2001. ‘Biofilms and device-associated infections’, Emerging infectious diseases, 7: 277.

Dunne, W. M. 1990. ‘Effects of subinhibitory concentrations of vancomycin or cefamandole on biofilm production by coagulase-negative staphylococci’, Antimicrobial Agents and Chemotherapy, 34: 390.

Elbing, Karen, and Roger Brent. 2002. ‘Media Preparation and Bacteriological Tools’, Current Protocols in Molecular Biology, 59: 1.1.1–1.1.7.

Elemam, Azza, Joseph Rahimian, and William Mandell. 2009. ‘Infection with panresistant Klebsiella pneumoniae: a report of 2 cases and a brief review of the literature’, Clinical infectious diseases, 49: 271–74.

Geredew Kifelew, Legesse, James G. Mitchell, and Peter Speck. 2019. ‘Mini-review: efficacy of lytic bacteriophages on multispecies biofilms’, Biofouling, 35: 472–81.

Górski, Andrzej, and Beata Weber-Dabrowska. 2005. ‘The potential role of endogenous bacteriophages in controlling invading pathogens’, Cellular and molecular life sciences, 62: 511.

Hamilton-Miller, J. M. T. 1965. ‘Modes of resistance to benzylpenicillin and ampicillin in twelve Klebsiella strains’, Microbiology, 41: 175–84.

Hennequin, Claire, Claire Aumeran, Frédéric Robin, Ousmane Traore, and Christiane Forestier. 2012. ‘Antibiotic resistance and plasmid transfer capacity in biofilm formed with a CTX-M-15-producing Klebsiella pneumoniae isolate’, Journal of Antimicrobial Chemotherapy, 67: 2123–30.

Hidron, Alicia I., Jonathan R. Edwards, Jean Patel, Teresa C. Horan, Dawn M. Sievert, Daniel A. Pollock, and Scott K. Fridkin. 2008. ‘Antimicrobial-resistant pathogens associated with healthcare-associated infections: annual summary of data reported to the National Healthcare Safety Network at the Centers for Disease Control and Prevention, 2006–2007’, Infection Control & Hospital Epidemiology, 29: 996–1011.

Huang, Su-Hua, Chien-Kuo Wang, Hwei-Ling Peng, Chien-Chen Wu, Ying-Tsong Chen, Yi-Ming Hong, and Ching-Ting Lin. 2012. ‘Role of the small RNA RyhB in the Fur regulon in mediating the capsular polysaccharide biosynthesis and iron acquisition systems in Klebsiella pneumoniae’, BMC Microbiology, 12: 148.

Hughes, Kevin A., Ian W. Sutherland, and Martin V. Jones. 1998. ‘Biofilm susceptibility to bacteriophage attack: the role of phage-borne polysaccharide depolymerase’, Microbiology, 144: 3039–47.

Jamala, Muhsin, Tahir Hussaina, C. Dasb, and Saadia Andleeba. 2015. ‘Inhibition of clinical multi-drug resistant Klebsiella pneumoniae biofilm by Siphoviridae bacteriophage Z’, Science Letters, 3: 122–6.

Kirchner, Sebastian, Joanne L. Fothergill, Elli A. Wright, Chloe E. James, Eilidh Mowat, and Craig Winstanley. 2012. ‘Use of artificial sputum medium to test antibiotic efficacy against Pseudomonas aeruginosa in conditions more relevant to the cystic fibrosis lung’, Journal of visualized experiments : JoVE: e3857–e57.

Kutateladze, Mzia, and Revaz Adamia. 2010. ‘Bacteriophages as potential new therapeutics to replace or supplement antibiotics’, Trends in Biotechnology, 28: 591–95.

Lehman, Susan M., and Rodney M. Donlan. 2015. ‘Bacteriophage-mediated control of a two-species biofilm formed by microorganisms causing catheter-associated urinary tract infections in an in vitro urinary catheter model’, Antimicrobial agents and chemotherapy, 59: 1127–37.

Liao, K. S., S. M. Lehman, D. J. Tweardy, R. M. Donlan, and B. W. Trautner. 2012. ‘Bacteriophages are synergistic with bacterial interference for the prevention of Pseudomonas aeruginosa biofilm formation on urinary catheters’, Journal of applied microbiology, 113: 1530–39.

Majkowska-Skrobek, Grazyna, Agnieszka Latka, Rita Berisio, Barbara Maciejewska, Flavia Squeglia, Maria Romano, Rob Lavigne, Carsten Struve, and Zuzanna Drulis-Kawa. 2016. ‘Capsule-targeting depolymerase, derived from Klebsiella KP36 phage, as a tool for the development of anti-virulent strategy’, Viruses, 8: 324.

Milo, Scarlet, Hollie Hathaway, Jonathan Nzakizwanayo, Diana R. Alves, Patricia Pérez Esteban, Brian V. Jones, and A. Toby A. Jenkins. 2017. ‘Prevention of encrustation and blockage of urinary catheters by Proteus mirabilis via pH-triggered release of bacteriophage’, Journal of Materials Chemistry B, 5: 5403–11.

NICE. 2018. “Urinary tract infection (catheter-associated): antimicrobial prescribing.” In NICE guideline [NG113]. The National Institute for Health and Care Excellence.

Nzakizwanayo, Jonathan, Aurélie Hanin, Diana R. Alves, Benjamin McCutcheon, Cinzia Dedi, Jonathan Salvage, Karen Knox, Bruce Stewart, Anthony Metcalfe, and Jason Clark. 2016. ‘Bacteriophage can prevent encrustation and blockage of urinary catheters by Proteus mirabilis’, Antimicrobial agents and chemotherapy, 60: 1530–36.

Parisien, A, B Allain, J Zhang, R Mandeville, and CQ Lan. 2008. ‘Novel alternatives to antibiotics: bacteriophages, bacterial cell wall hydrolases, and antimicrobial peptides’, Journal of Applied Microbiology, 104: 1–13.

Podschun, Rainer, and U Ullmann. 1998. ‘Klebsiella spp. as nosocomial pathogens: epidemiology, taxonomy, typing methods, and pathogenicity factors’, Clinical microbiology reviews, 11: 589–603.

Ratner, Hannah K., Timothy R. Sampson, and David S. Weiss. 2015. ‘I can see CRISPR now, even when phage are gone: a view on alternative CRISPR-Cas functions from the prokaryotic envelope’, Current opinion in infectious diseases, 28: 267–74.

Ryan, Elizabeth M., Sean P. Gorman, Ryan F. Donnelly, and Brendan F. Gilmore. 2011. ‘Recent advances in bacteriophage therapy: how delivery routes, formulation, concentration and timing influence the success of phage therapy’, Journal of Pharmacy and Pharmacology, 63: 1253–64.

Sabir, Nargis, Aamer Ikram, Gohar Zaman, Luqman Satti, Adeel Gardezi, Abeera Ahmed, and Parvez Ahmed. 2017. ‘Bacterial biofilm-based catheter-associated urinary tract infections: causative pathogens and antibiotic resistance’, American journal of infection control, 45: 1101–05.

Sanchez, Guillermo V, Ronald N Master, Richard B Clark, Madiha Fyyaz, Padmaraj Duvvuri, Gupta Ekta, and Jose Bordon. 2013. ‘Klebsiella pneumoniae antimicrobial drug resistance, United States, 1998–2010’, Emerging infectious diseases, 19: 133.

Singla, Saloni, Kusum Harjai, and Sanjay Chhibber. 2014. ‘Artificial Klebsiella pneumoniae biofilm model mimicking in vivo system: altered morphological characteristics and antibiotic resistance’, The Journal of Antibiotics, 67: 305–09.

Stewart, M. H., and B. H. Olson. 1992. ‘Physiological studies of chloramine resistance developed by Klebsiella pneumoniae under low-nutrient growth conditions’, Applied and Environmental Microbiology, 58: 2918.

Storms, Zachary J., Matthew R. Teel, Kevin Mercurio, and Dominic Sauvageau. 2019. ‘The Virulence Index: A Metric for Quantitative Analysis of Phage Virulence’, PHAGE, 1: 17–26.

Tacconelli, Evelina, Elena Carrara, Alessia Savoldi, Stephan Harbarth, Marc Mendelson, Dominique L Monnet, Céline Pulcini, Gunnar Kahlmeter, Jan Kluytmans, and Yehuda Carmeli. 2018. ‘Discovery, research, and development of new antibiotics: the WHO priority list of antibiotic-resistant bacteria and tuberculosis’, The Lancet Infectious Diseases, 18: 318–27.

Tait, K., L. C. Skillman, and I. W. Sutherland. 2002. ‘The efficacy of bacteriophage as a method of biofilm eradication’, Biofouling, 18: 305–11.

Townsend, Eleanor, Lucy Kelly, Lucy Gannon, George Muscatt, Rhys Dunstan, Slawomir Michniewski, Hari Sapkota, Saija J Kiljunen, Anna Kolsi, Mikael Skurnik, Trevor Lithgow, Andrew D MIllard and Eleanor Jameson. 2020. ‘Isolation and chacterisation of Klebsiella phages for phage therapy’, bioRxiv: 2020.07.05.179689.

Ujmajuridze, Aleksandre, Nina Chanishvili, Marina Goderdzishvili, Lorenz Leitner, Ulrich Mehnert, Archil Chkhotua, Thomas M. Kessler, and Wilbert Sybesma. 2018. ‘Adapted Bacteriophages for Treating Urinary Tract Infections’, Frontiers in Microbiology, 9: 1832.

Van Laar, Tricia A, Tsute Chen, Tao You, and Kai P Leung. 2015. ‘Sublethal concentrations of carbapenems alter cell morphology and genomic expression of Klebsiella pneumoniae biofilms’, Antimicrobial Agents and Chemotherapy, 59: 1707–17.

Verma, Vivek, Kusum Harjai, and Sanjay Chhibber. 2010. ‘Structural changes induced by a lytic bacteriophage make ciprofloxacin effective against older biofilm of Klebsiella pneumoniae’, Biofouling, 26: 729–37.

Wazait, H. D., H. R. H. Patel, V. Veer, M. Kelsey, J. H. P. Van Der Meulen, R. A. Miller, and M. Emberton. 2003. ‘Catheter-associated urinary tract infections: prevalence of uropathogens and pattern of antimicrobial resistance in a UK hospital (1996–2001)’, BJU international, 91: 806–09.

Wu, Yunqiang, Rui Wang, Mengsha Xu, Yanan Liu, Xianchao Zhu, Jiangfeng Qiu, Qiming Liu, Ping He, and Qingtian Li. 2019. ‘A Novel Polysaccharide Depolymerase Encoded by the Phage SH-KP152226 Confers Specific Activity Against Multidrug-Resistant Klebsiella pneumoniae via Biofilm Degradation’, Frontiers in Microbiology, 10: 2768.

Yang, D, and Z Zhang. 2008. ‘Biofilm-forming Klebsiella pneumoniae strains have greater likelihood of producing extended-spectrum β-lactamases’, Journal of Hospital Infection, 68: 369–71.

Zahller, Jeff, and Philip S Stewart. 2002. ‘Transmission electron microscopic study of antibiotic action on Klebsiella pneumoniae biofilm’, Antimicrobial Agents and Chemotherapy, 46: 2679–83.

